# Reflectins form multicompartment liquid-liquid phase separated condensates that mirror and may facilitate spatial organization in squid skin Bragg lamellae

**DOI:** 10.64898/2026.03.13.711711

**Authors:** Reid Gordon, Robert Levenson, Brandon Malady, Yahya Al Sabeh, Daniel E. Morse

## Abstract

Cationic reflectin proteins transduce neuronal signals tuning skin color for dynamic camouflage and communication in Loliginid squid. Neuronally released acetylcholine (ACh) activates phosphorylation of the reflectins, triggering their condensation, folding and hierarchical assembly. This causes osmotic and Gibbs-Donnan dehydration of membrane-enclosed Bragg lamellae containing these proteins in skin cells called iridocytes, changing their refractive index and spacing to finely tune the wavelength of reflected light. Reflectins B and C are enriched in ACh-responsive iridocytes, suggesting they are critical for the neuronally tuned protein phase transitions within the Bragg lamellae. Here, using pH titration as an *in vitro* surrogate for phosphorylation to reduce protein net charge density, we demonstrate with confocal microscopy that reflectins A1, A2, B and C individually undergo LLPS to form protein dense liquid condensates, while physiological mixtures of these proteins exhibit coordinated phase transitions to form multicompartment condensates whose internal spatial organization is tuned by the proportions and charge densities of the different reflectins. Our findings demonstrate that (i) the charge densities of the different reflectins control their spatial organization within liquid condensates, (ii) the relative proportions of the different reflectins found in the ACh-responsive iridocytes increase the sensitivity of their proteins’ liquid phase transitions to changes in reflectin charge density; and (iii) the spatial segregation of reflectins A1 and C we observe in their multiphasic liquid condensate mirror their spatial segregation in the Bragg lamellae *in vivo*, suggesting a mechanistic explanation for the spatial segregation of these reflectins across the iridocytes’ Bragg lamellae.

## Introduction

The striking dynamic coloration cephalopods use for communication and camouflage is enabled by a complex optical system that combines pigmentary and structural coloration (1–4). Leucophores scatter broadband light through Mie scattering and iridocytes reflect light in a wavelength- and angle-dependent manner via regular and extensive membrane invaginations which form stacks of Bragg lamellae densely filled with reflectin proteins (5–8). Uniquely, *Loliginidae* squid can neuronally control the reflectance of leucophores and iridocytes (9, 10). Activation of iridocytes by neuronally released acetylcholine by (11) causes the phosphorylation of cationic reflectin proteins (5, 6, 12). This decreases Coulombic repulsion and progressively drives the reversible assembly and condensation of reflectin proteins (13, 14), with the resulting reversible osmotic and Gibbs-Donnan dehydration of the Bragg lamellae (15). This reversible and proportional dehydration reduces the thickness and spacing of the Bragg lamellae and increases their refractive index, thus decreasing the wavelength and increasing the intensity of reflected light (7, 8, 15). A similar reversible and tunable mechanism controls the dehydration and resulting broad-band reflectivity of reflectin-enriched vesicles in leucophores (16, 17).

Proteins in the reflectin family typically contain a unique methionine-rich conserved N-terminal motif (RMn) and are block-copolymers of conserved reflectin repeat motifs (RMs) separated by cation linker regions (**Figure 1A,B**) (13, 18, 19). They also are enriched in arginine, histidine and aromatic residues and contain little if any lysine and aliphatic residues (6, 12, 19). Reflectin A1 from *Doryteuthis opalascens* is an intrinsically disordered monomer in acidic conditions, while transitioning to extensive secondary structure as detected by circular dichroism in neutral conditions (20). As a surrogate for *in vivo* reflectin phosphorylation, reduction of protein net charge density (NCD) by progressive pH titration of histidine residues, phosphomimetic mutation by genetic insertion of negatively charged glutamate residues at known *in vivo* phosphorylation sites, or low-voltage electroreduction drives reflectin folding and assembly as monitored by dynamic light scattering (DLS), TEM and circular dichroism (CD) (13, 20–22). This reversible folding and assembly is opposed by Coulombic repulsion of the cationic linker regions, with the protein’s NCD precisely controlling folding and assembly size in low-salt conditions (13, 20). Increasing salt concentration drives reflectin A1 assembly by electrostatic screening (14, 23) and by increasing the contribution of the hydrophobic effect ultimately leading to liquid-liquid phase separation (23).

**Figure 1.**
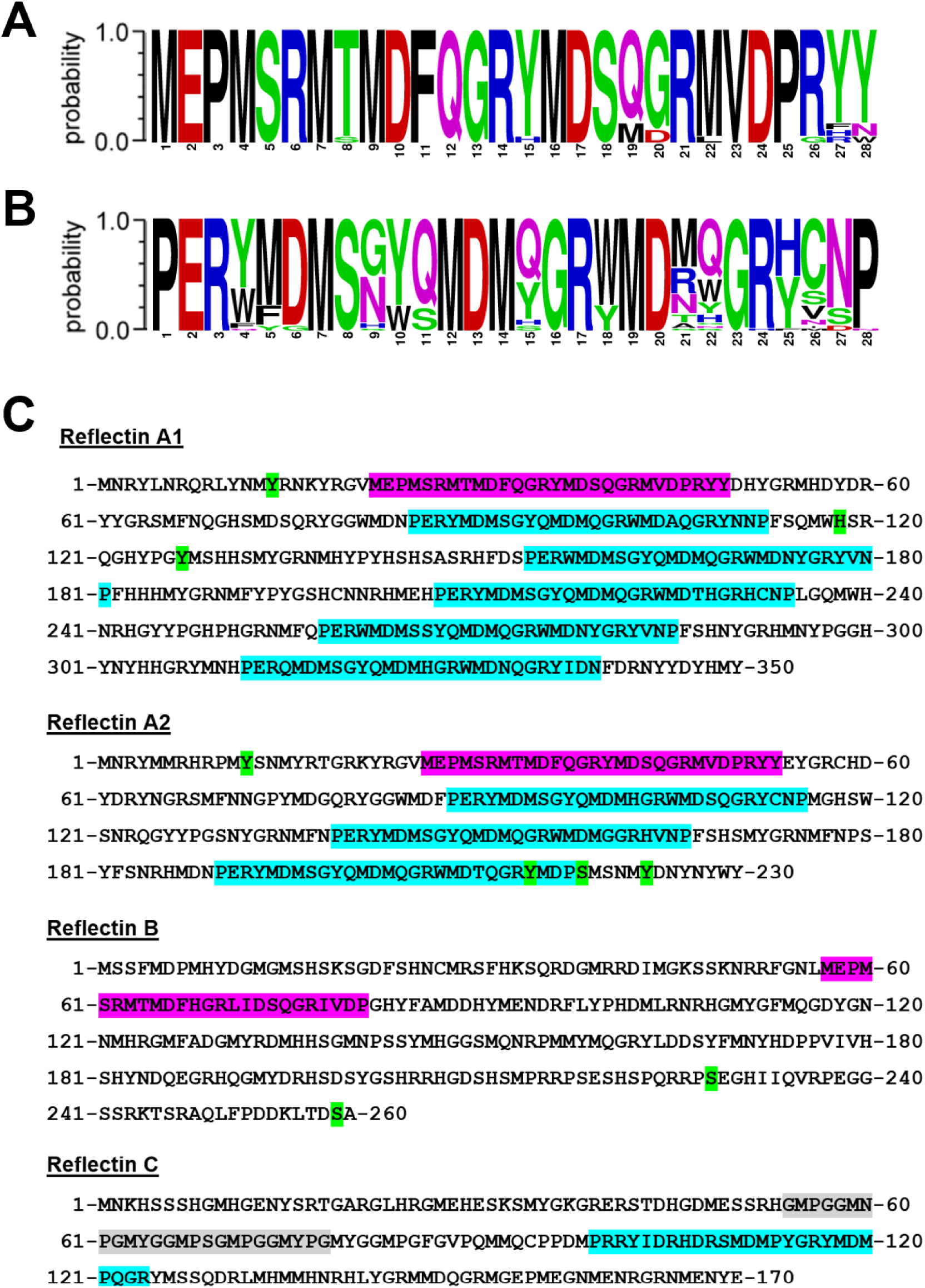
Reflectin proteins contain strongly conserved motifs. Sequence logos from the alignment of the (A) N-terminal and (B) reflectin repeat motifs from 51 reflectin proteins found in *Octopus bimaculoides, Euprymna scolopes, Sepia officinalis, Doryteuthis opalascens* and *Doryteuthis pealeii.* C) Sequences of reflectins A1, A2, B, and C from Doryteuthis opalascens. N-terminal motifs are highlighted in magenta and repeat motifs in teal. Reflectin C contains one poorly conserved repeat motif (red) and the GMXX proline-rich region (gray). Green residues are phosphorylated (reflectins A1, A2) or dephosphorylated (reflectin B) upon iridocyte activation by acetylcholine. Sequence logo created using U.C. Berkeley WebLogo.

Protein liquid-liquid phase separation is a thermodynamic process in which a single-phase protein solution separates into coexisting protein-dilute and protein-dense phases in equilibrium (24, 25). *In vivo*, phase separation is commonly regulated by phosphorylation of the protein constituents (26, 27); the resulting transient liquid-like protein-dense condensates may exhibit dynamic material properties and, in the case of the nucleolus, form multicompartment condensates that vectorially organize ribosome biogenesis (28, 29). The observed *in vitro* protein NCD-dependent transition of reflectin A1 from discrete 10-100 nm diam. assemblies to liquid condensates mirrors *in* vivo observations of ultrastructural changes that demonstrate liquid-liquid phase separation in tunable iridocyte Bragg lamellae and leucophore vesicles upon acetylcholine activation (5, 9, 17). However, the most abundant protein components of tunable Bragg lamellae are reflectins B and C (6), and beyond other *Loliginid* squids (that also contain reflectin B) and *Doryteuthis opalascens* (that uniquely contains reflectin C), no closely homologous proteins have been described. Reflectin B contains the n-terminal RMn but lacks the repeated RMs; reflectin C contains a unique proline-rich region that is predicted to be disordered **(Figure S1)**, lacks the RMn and has one poorly conserved RM (18). Reflectins B and C are enriched in the squid’s dorsal, neurotransmitter-responsive iridocytes relative to ventral, non-responsive iridocytes from *Doryteuthis opalascens* (6). They are spatially organized within the lamellae, with reflectins A2, A2 and B localized to the center of the lamellae and reflectin C localized to the edge of the lamellar membrane (6). Like reflectin A1, reflectins A2, B and C also form soluble protein assemblies and pH titration demonstrated these assembly sizes are similarly controlled by protein NCD (13). A mixture of reflectins A1, A2, B and C in the same molar ratio as that found in neurotransmitter-responsive iridocytes also formed low polydispersity oligomers of protein NCD-dependent sizes.

This led us to hypothesize that these proteins could facilitate the previously observed ACh-triggered dehydration of the Bragg lamellae (6), and thus enhance the tunability of the reflected wavelength, by biophysically altering phase changes and liquid-like condensate properties in the Bragg lamellae. Here, we demonstrate using confocal microscopy that reflectins A2, B and C individually undergo liquid-liquid phase separation to form liquid-like condensates in a salt concentration- and protein NCD-dependent manner. Mixtures of reflectins A2, B, or C with reflectin A1 in the ratios found in neurotransmitter-responsive iridocytes show coordinated phase behavior, in which reflectins B and C sharpen the liquid phase boundary of A1 as a function of protein NCD. Further, at increased protein concentrations reflectins A1, A2, B and C form multicompartment liquid-like condensates whose spatial organization is tuned by protein NCD and proportions of reflectin protein species, thus recapitulating the most striking observation of *in vivo* segregation in the Bragg lamellae. Liquid condensates formed by reflectins B or C show greater diffusivity than those of A1, and reflectin C increases the diffusivity of reflectin A1 in A1/C condensates. These results introduce novel micrometer-scale spatial tunability for reflectin-based biomaterials and indicate specific roles through which the complex interplay between these four proteins could enhance the protein NCD-triggered dehydration of Bragg lamellae.

## Results

### Reflectins A1, A2, B and C each undergo LLPS to form liquid-like condensates

To test the hypothesis that previously observed coordinated assembly of reflectins A1, A2, B and C (13) may extend to coordinated phase separation of the four reflectin proteins, we analyzed the liquid phase boundaries of A1(23) and in this study, of A2, B and C as well as physiological mixtures of A1, A2, B and C as a function of NaCl concentration and protein NCD by pH titration. The liquid phase boundary of reflectin A2 for pH values 4 to 7.5 was determined by dilution of protein stock into respective buffers of varying ionic strength as previously performed for reflectin A1 (23). Liquid-like condensates (**Figure 2A**) were distinguished from protein assemblies (**Figure 2B**) (13) by their significantly larger size and ability to fuse and relax to sphericity **(Figure S1A)** (23, 24). The dilution of protein stock containing reflectins A1 and A2 in the ratio found in neurotransmitter-tunable iridocytes resulted in co-phase separation of both proteins into liquid condensates (**Figure 2C**). For all conditions tested, this mixture displayed a coordinated phase transition: either assemblies or liquid condensates containing both proteins were observed (**Figure 2C,D**), while a solution containing liquid condensates of one protein and assemblies of the other was never observed. Of reflectins A2, B and C, A2 is most similar to A1 regarding the protein sequences and block copolymer organization of RMs and cationic linkers (**Figure 1, 2F)**. The liquid phase boundaries of A2 and A1/A2 mixture both display similar inverse relationships between protein NCD and the NaCl concentration needed to drive LLPS, and the slope of this relationship is approximately constant over the range of protein NCDs tested (**Figure 2E, S2)**. These results match those from previous analyses of reflectin A1 and A2 assembly sizes as a function of protein NCD (20). Interestingly, at protein NCDs less than 9 the mixture of A1/A2 had a lower phase boundary as a function of NaCl concentration than either A1 or A2 alone, showing that interactions between the two proteins encourages LLPS at low protein NCDs (**Figure 2E**).

**Figure 2.**
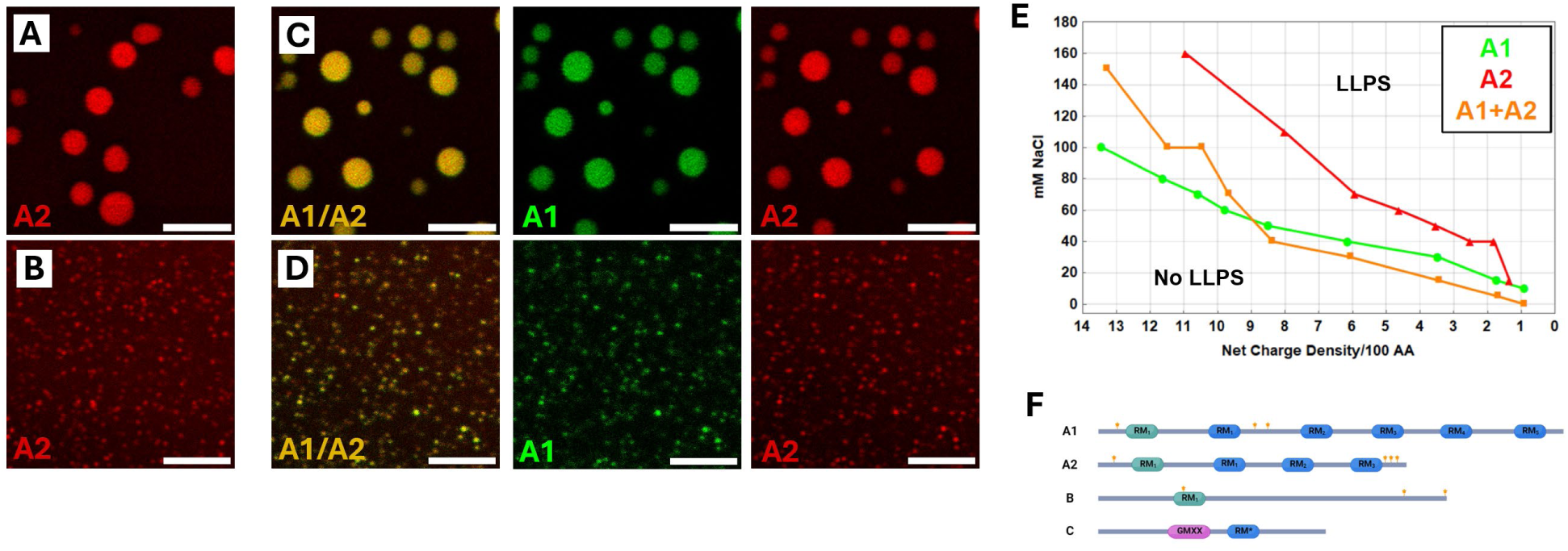
Liquid phase boundary of reflectin A2 and A1/A2 determined by confocal microscopy. One hundred μM partially fluorescently labeled reflectin A2 (see methods: protein labeling) or 52.6 and 47.37 μM of partially fluorescently labeled A1 and A2 respectively was diluted to a final protein concentration of 4 μM into buffers ranging from pH 4 to 8 and NaCl concentrations from 0 to 300 mM (See methods: phase diagram). (A) Liquid-like protein-dense condensates and (B) assemblies of reflectin A2. C) Liquid-like protein-dense condensates and (D) assemblies of reflectin A1 and A2. E) Liquid phase boundary as a function of NaCl concentration and calculated protein NCD of reflectins A1(green), A2 (red) and A1/A2 (orange). F) Representations of the sequences of reflectin A1 and A2 showing the RMn (green), RMs (blue), cationic linkers (grey), and known sites of *in vivo* phosphorylation (stars). Protein net charge density was calculated using pKas of the amino acid residues, and A1/A2 liquid phase boundary is shown using the protein NCDs of A1. Reflectin A1-only data are from (23). All scale bars are 5 μm.

Identical analysis of the liquid phase boundary of reflectin B showed that the protein can form liquid condensates (**Figure 3A**), assemblies (**Figure 3B**), or arrested gel-like clusters (**Figure 3C**) which did not display the fusion and relaxation to sphericity characteristic of the liquid-like condensates **(Figure S1B)**. For mixtures of reflectins A1 and B, the phase transitions of both proteins to form liquid condensates (**Figure 3D**), assemblies (**Figure 3E**) or gel like clusters (**Figure 3F**) were coordinated for all observations. The slope of the liquid phase boundary of reflectin B is similar to that of reflectin A1 except for protein NCDs greater than 9, with arrested gel-like clusters observed at protein NCDs of 11-14 (**Figure 3G, S3)**. Notably, as a function of NaCl concentration the liquid phase boundary of the reflectin A1/B mixture is lower than that of reflectin A1 for protein NCDs less than approximately 4 and higher for protein NCDs greater than approximately 8, demonstrating that reflectin B can both promote or restrict LLPS of reflectin A1 (**Figure 3G, S3)**. Reflectin C formed liquid-like condensates (**Figure 4A**) or assemblies (**Figure 4B**) in a protein NCD and NaCl concentration-dependent manner (**Figure 4E, S4)** and these condensates displayed fusion and relaxation to sphericity **(Figure S1)**. As with reflectins A2 and B, reflectins A1 and C co-phase separated to form liquid condensates containing both proteins (**Figure 4C**) or assemblies containing both proteins (**Figure 4D**), and these transitions were coordinated. As a function of ionic strength, for spaces in the phase diagram above the liquid phase boundary of reflectin A1 but below that of reflectin C with (**Figure 4E**), a mixture of reflectin A1/C never displayed the co-existence of reflectin A1 condensates and reflectin C assemblies. Reflectin C, which lacks the RMn and has only one poorly conserved RM (**Figure 1**), had the greatest effect of inhibiting the LLPS of reflectin A1 at high and intermediate protein NCDs (**Figure 4E**).

**Figure 3.**
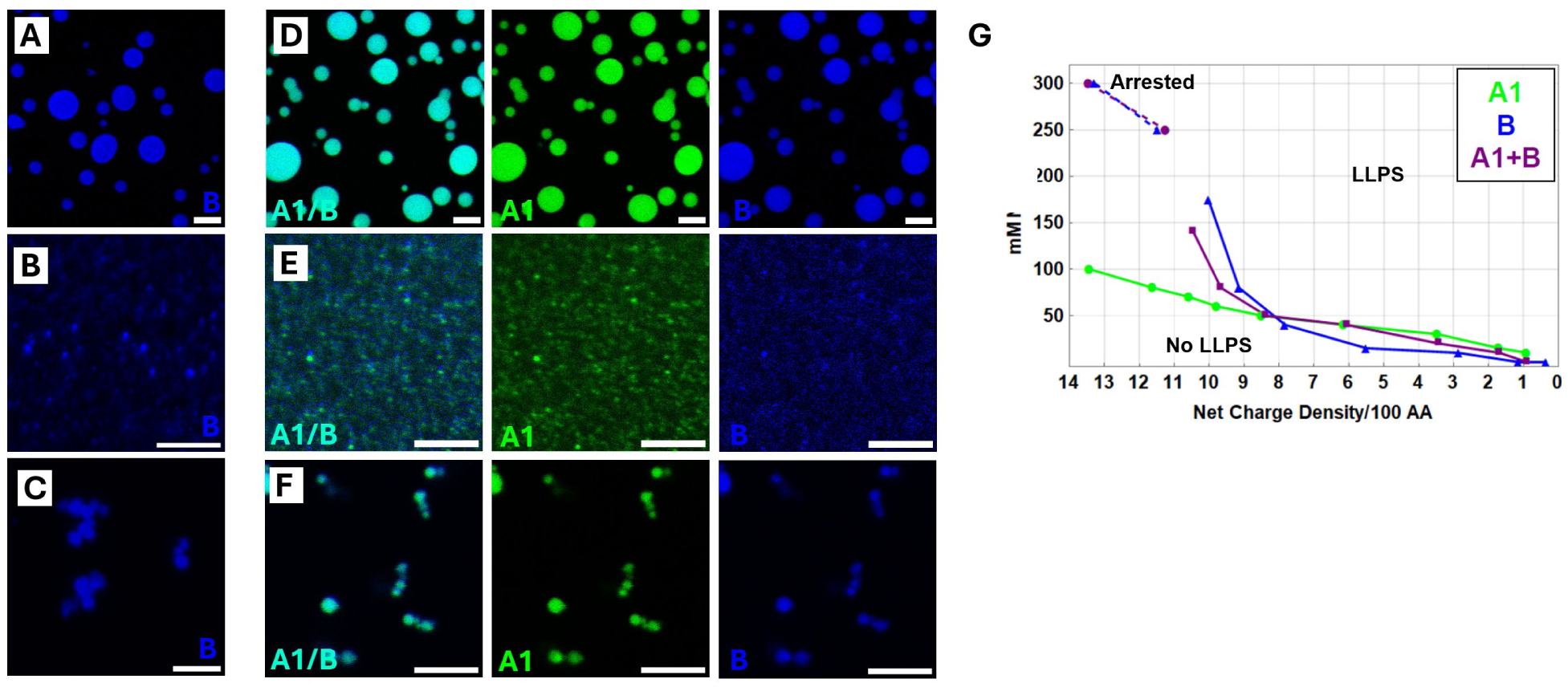
Liquid phase boundary of reflectin B and A1/B determined by confocal microscopy. One hundred μM partially fluorescently labeled reflectin B or a solution of 24.4 and 75.6 μM of partially fluorescently labeled A1 and B respectively was diluted to a final protein concentration of 4 μM into buffers ranging from pH 4 to 8 and NaCl concentrations from 0 to 300 mM. A) Liquid-like protein-dense condensates, (B) assemblies and (C) arrested clusters of reflectin B. D) Liquid-like protein-dense condensates, (E) assemblies and (F) arrested clusters of reflectin A1 and B. G) Liquid phase boundary as a function of NaCl concentration and calculated protein NCD of reflectins A1 (green), B (blue) and A1/B (purple). Dashed line represents boundary between assemblies and arrested clusters. A1/B liquid phase boundary is shown using the protein NCDs of A1. All scale bars are 5 μm.

**Figure 4.**
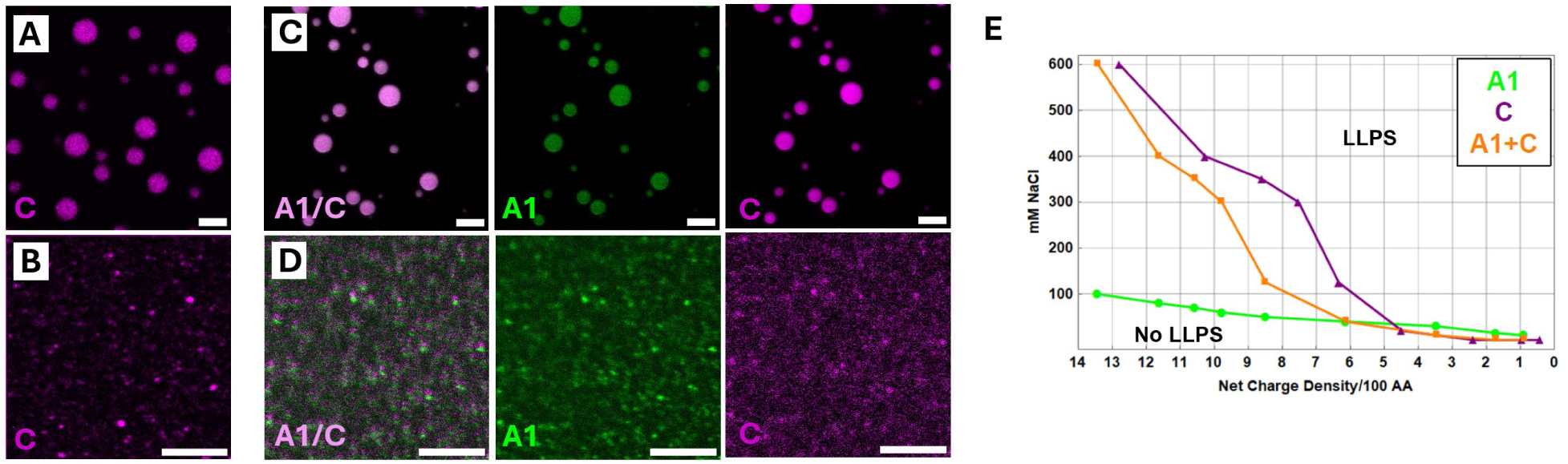
Liquid phase boundary of reflectin C and A1/C determined by confocal microscopy. One hundred μM partially fluorescently labeled reflectin C or a solution of 21.7 and 78.6 μM of partially fluorescently labeled A1 and C respectively was diluted to a final protein concentration of 4 μM into buffers ranging from pH 4 to 8 and NaCl concentrations from 0 to 300 mM. A) Liquid-like protein-dense condensates and (B) assemblies of reflectin C. C) Liquid-like protein-dense condensates and (D) assemblies (bottom) of reflectins A1 and C. A1/C liquid phase boundary is shown using the protein NCDs of A1. All scale bars are 5 μm. E) Liquid phase boundary as a function of NaCl concentration and calculated protein NCD of reflectins A1 (green), C (purple) and A1/C (orange).

After observing the pronounced changes of the liquid phase boundary of A1 by A2, B and C, we hypothesized that the different ratios of these four reflectins in neurotransmitter-tunable and non-tunable iridocytes in *Doryteuthis opalascens* (**Figure 5G**) (6) might sensitize the response of the *in vivo* phase changes to decreasing protein NCD induced by phosphorylation. We investigated the phase behavior of mixtures of reflectins A1, A2, B and C at 4 μM total protein concentration and found the phase transition from soluble assemblies to two separate liquid phases of all four reflectins was coordinated (**Figure 5A,B,C,D,E,F)** as was observed with binary mixtures of A1/B, A1/A2 and A1/C. We then compared the phase diagrams of reflectins A1, A2, B and C (**Figure 5H**) in ratios characteristic of the neurotransmitter-responsive and -nonresponsive cells, as a function of protein NCD and NaCl concentration. Surprisingly, these two liquid phase boundaries were nearly identical to one another, and to that of reflectin B alone (**Figure 3G, 5H, S5)**, demonstrating that the lower proportion of reflectin B in the nonresponsive mixture of A1, A2, B and C is sufficient to control the liquid phase boundaries of the other three proteins. It is notable that A2, B and C all significantly inhibit LLPS of A1 at high protein NCDS despite these proteins having lower protein NCDs than A1 at each pH. We conclude that these proteins inhibit LLPS of A1 at high protein NCDs through a mechanism different from altering the overall protein NCD of the A1/A2, A1/B, A1/C and A1/A2/B/C mixtures, and different from simple dilution of the total protein concentration of A1 **(Figure S6)**.

**Figure 5.**
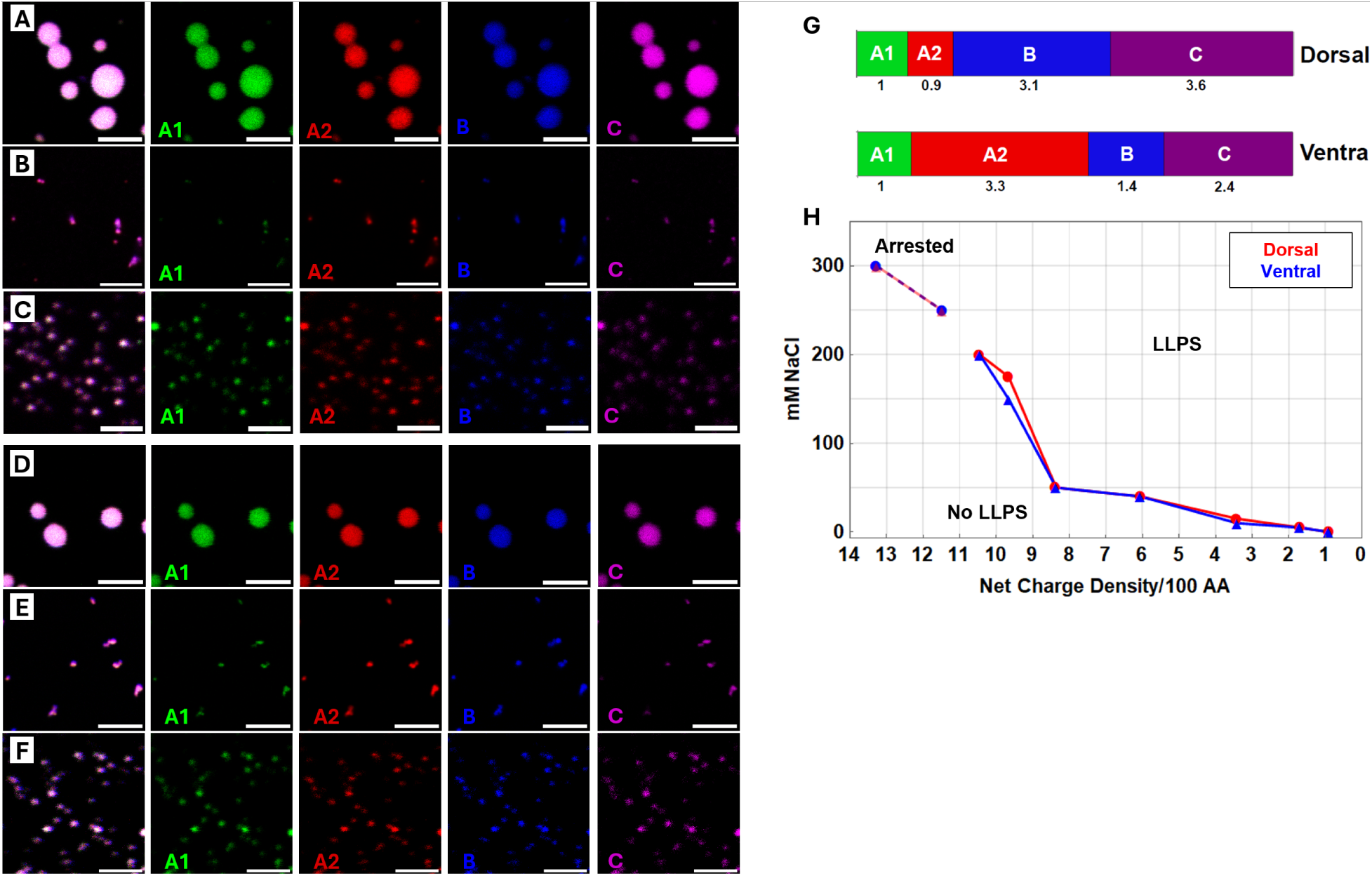
Liquid phase boundary of mixtures of A1, A2, B and C found in neurotransmitter-tunable and non-tunable iridocytes determined by confocal microscopy. Mixtures comprising total protein concentrations of 100 μM reflectins A1, A2, B and C in the molar ratios found in either the ACh-responsive or nonresponsive iridocytes were diluted to a final protein concentration of 4 μM into buffers ranging from pH 4 to 8 and NaCl concentrations from 0 to 300 mM. A) Liquid-like protein-dense condensates B) arrested clusters and C) assemblies formed by the ACh-responsive mixture of A1, A2, B and C. D) Liquid-like protein-dense condensates, E) arrested clusters and F) assemblies formed by ACh-nonresponsive mixture of A1, A2, B and C. G) Previously determined mole percent of reflectins found in ACh-responsive and nonresponsive iridocyte cells from *Doryteuthis opalescens* normalized to their concentrations of A1(6). H) Liquid phase boundary (solid line) and assembly/cluster boundary (dashed line) of ACh-responsive and nonresponsive mixtures as a function of calculated protein NCD of A1 and NaCl concentration. All scale bars are 5 μm.

### Multiphase condensate organization determined by protein composition and NCD

To test the hypothesis that condensate multiphase organization is dynamically tuned by protein NCD, we examined multiphase organization as a function of protein NCD for both neurotransmitter-responsive and nonresponsive mixtures of reflectins A1, A2, B and C at a total protein concentration of 8 μM. At pH 5.5, the high protein NCD condition, all four proteins were evenly distributed throughout the condensates (**Figure 6A**). Reflectins A1 and A2 were not evenly distributed at pH 7 (lower protein NCD) and pH 8 (lowest protein NCD) where condensates exhibited nonuniform voids of these two proteins (**Figure 6B, C)**. In marked contrast, the responsive mixture of reflectins A1, A2, B and C at the lowest protein NCD condition formed condensates with A1 and A2 enriched in the periphery and shielding B and C from solvent (**Figure 6D**). Lowering net charge density rearranged this multiphase organization with reflectins A1 and A2 forming small distinct puncta in the condensate interior (**Figure 6 E, F)**. To determine the effect of individual reflectins on the organization of condensates formed by the responsive mixture, we compared condensates that lacked either reflectin A2, B or C to condensates containing all four proteins in responsive (**Figure 7A, B)** or non-responsive mixtures (**Figure 7C, D)**. Condensates lacking reflectin B (**Figure 7E, F**) displayed similar organization as those with B (**Figure 7A, B**). Condensates lacking reflectin A2 did not exhibit multiphase organization (**Figure 7G, H)**, suggesting that reflectin A2 in proportion found in the responsive mixture is necessary to drive multiphase organization. In condensates lacking reflectin C, reflectins A1 and A2 formed small compartments in the condensate interior (**Figure 7I, J)**. The segregation of reflectins A1 and A2 from reflectin B was the greatest and this was the only mixture that demonstrated the exclusion of reflectin B from the interior of the droplets, suggesting that reflectin C increases the solubility of reflectin B in reflectins A1 and A2.

**Figure 6.**
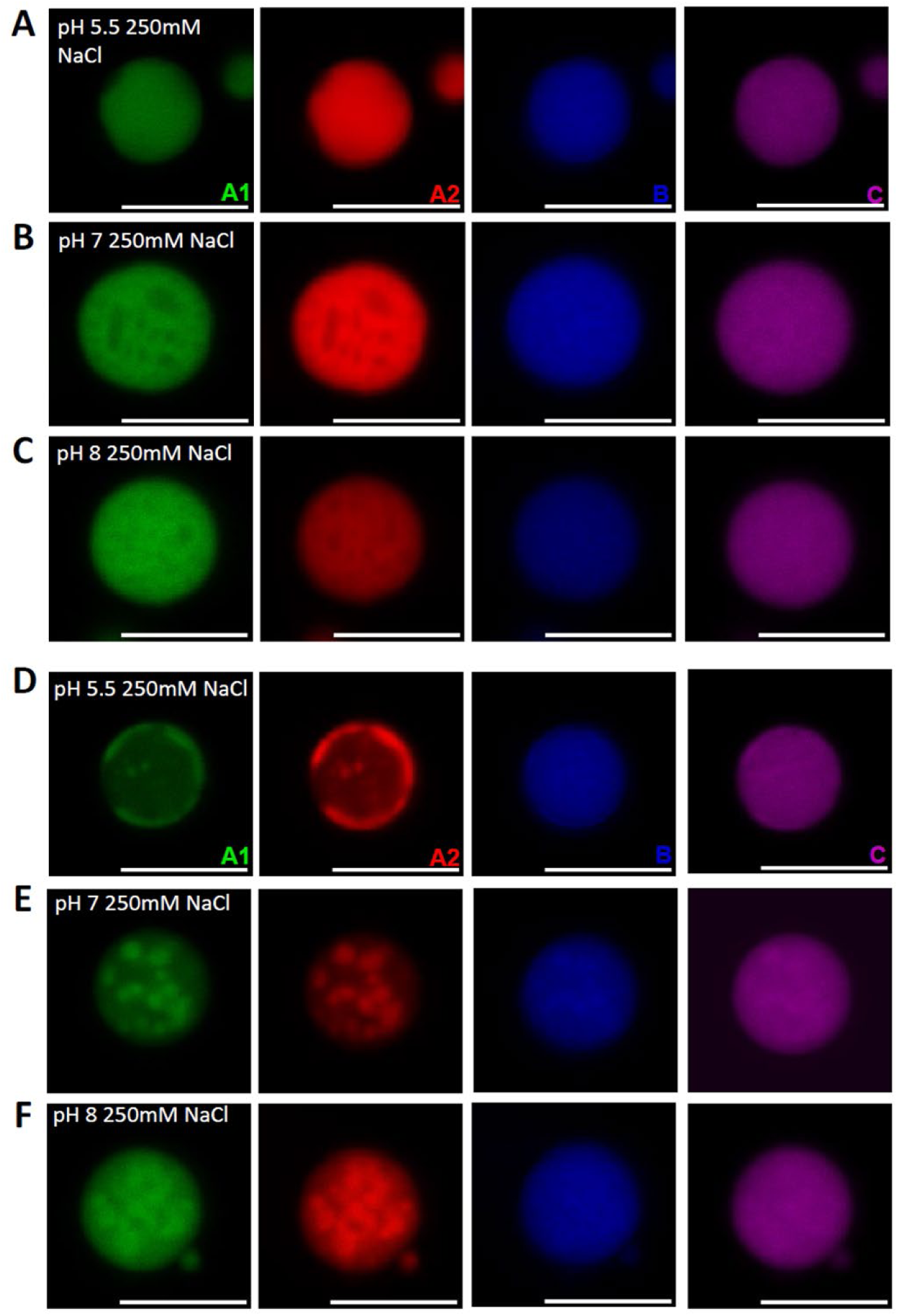
Single and multiphase condensates formed by neurotransmitter-tunable and non-tunable mixtures of A1, A2, B and **C.** Mixtures of 200 μM (total) reflectins A1, A2, B and C in the molar ratios found in either ACh-responsive or nonresponsive iridocytes were diluted to a final protein concentration of 8 μM into the indicated buffers. Fluorescence was imaged at the each of the wavelengths of emission of the uniquely labelled reflectins A1, A2, B and C in the ACh-nonresponsive mixture in 250 mM NaCl at (A) pH 5.5, (B) pH 7 and (C) pH 8. Similarly, fluorescence was imaged at the each of the wavelengths of emission of the uniquely labelled reflectins A1, A2, B and C in the ACh-responsive mixture in 250 mM NaCl at (D) pH 5.5, (E) pH 7 and (F) pH 8. All scale bars are 5 μm.

**Figure 7.**
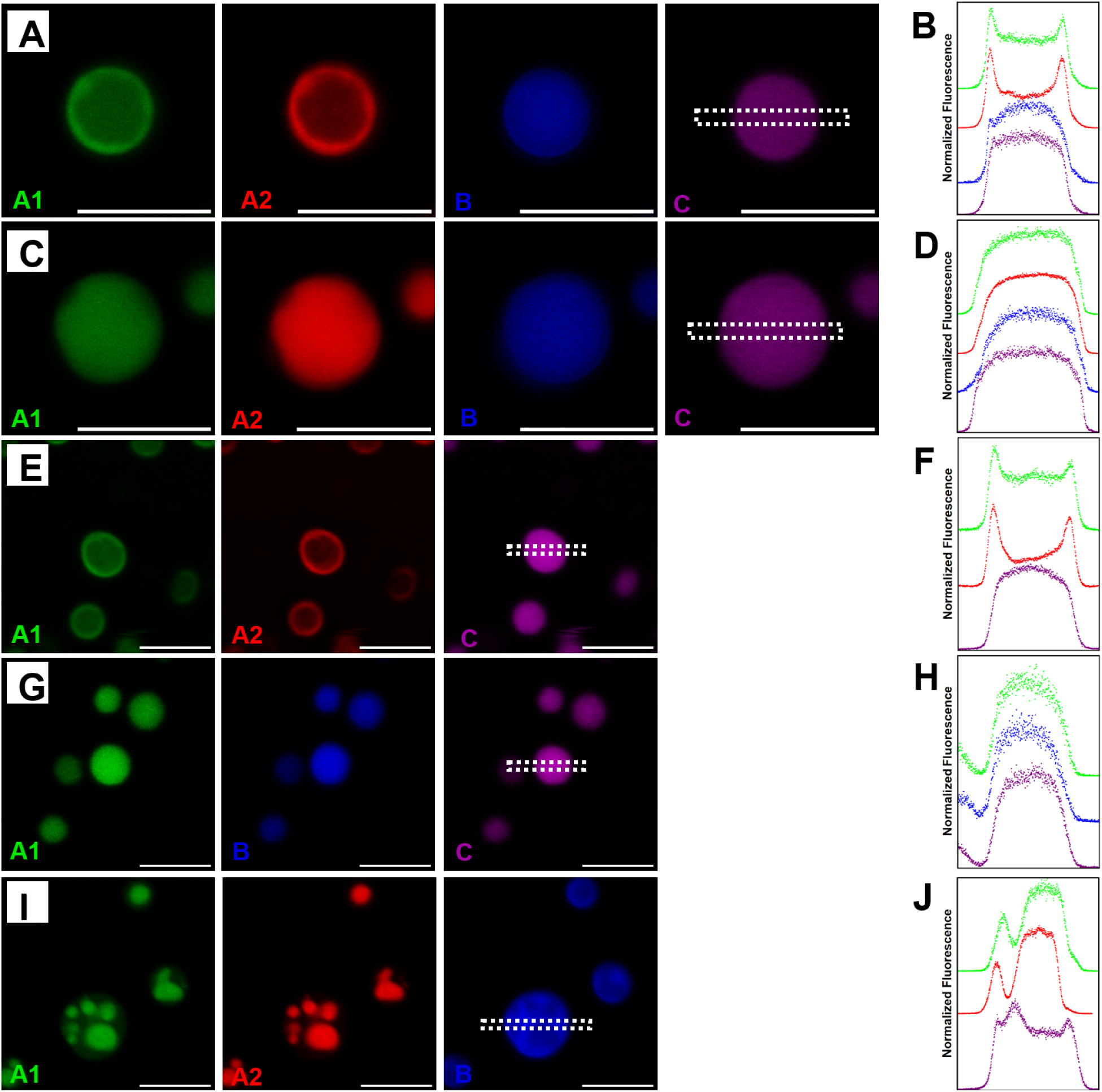
The effect of individual reflectins on the multiphase organization of condensates formed from neurotransmitter-tunable mixture of A1, A2, B and. **C.** Mixtures of 200 μM (total) of reflectins A1, A2, B and C found in either the ACh-responsive or ACh-nonresponsive iridocytes were diluted to a final protein concentration of 8 μM into into pH 5.5 (MES, 25 mM) buffer in 250 mM NaCl. (A) Fluorescence at the emission wavelengths of each of the uniquely labelled reflectins, and (B) fluorescence intensity across the dashed box for each of the images, for the ACh-responsive mixture. C) Fluorescence at the emission wavelengths of each of the uniquely labelled reflectins, and (D) fluorescence intensity across the dashed box for each of the images, for the ACh-nonresponsive mixture. (E) Individual channels and (F) fluorescence intensity across the dashed box for the ACh-responsive mixture of reflectins A1, B and C (omitting A2). (G) Individual channels and (H) fluorescence intensity across the dashed box for ACh-responsive mixture of reflectins A1, A2 and C (omitting B). (I) Individual channels and (J) fluorescence intensity across the dashed box for neurotransmitter-tunable mixture of reflectins A1, A2 and B (omitting C). All scale bars are 5 μm. (C) is reproduced from Figure 6A.

### Individual reflectins have different diffusivities within liquid-like condensates

To compare intra-condensate diffusivity we analyzed the fluorescence recovery after photobleaching (FRAP) of liquid-like condensates of reflectins B (**Figure 8A**) and C (**Figure 8B**). Fitting the resulting FRAP data (**Figure 8C**) to an equation for exponential decay revealed that the total fluorescence recovery for reflectin B condensates was 94.8 ± 2.5 % and for reflectin C condensates was 99.1 ±1.6 % (**Figure 8D**), demonstrating that all or nearly all of the dense phase protein is liquid-like in these condensates. The characteristic relaxation time for fluorescence recovery was 57.6 ± 12.6 s for reflectin B condensates and 12.2 ± 5.3 s for reflectin C condensates (**Figure 8E**). Previous *in vitro* FRAP analyses of reflectin A1 condensates at physiologically relevant pH values demonstrated relatively slow diffusivity and incomplete recovery (23). We investigated the effect of reflectin C on the diffusivity of reflectin A1 in condensates composed of both reflectin A1 and C by selectively photobleaching reflectin A1 and monitoring fluorescence recovery **(Figure 9 A, D)**. Plotting the characteristic relaxation time tau and fluorescence recovery against previously obtained data for condensates composed only of A1 (23) reveals that reflectin C dramatically increases the liquid-like population of reflectin A1 within A1/C condensates from 0.50 ± 0.04 to 0.93 ± 0.08 at pH 6 and from 0.32 ± 0.02 to 0.91 ± 0.02 at pH 7 **(Figure 9 B, E)**. Reflectin C similarly affected the rate of diffusion of A1 within A1/C condensates, decreasing tau from 641.57 ± 220.17 to 141.68 ± 26.26 s at pH 6 and from fluorescence recovery that was too slow to accurately fit to 182.64 ± 13.24 s at pH 7 **(Figure G C, F)**.

**Figure 8.**
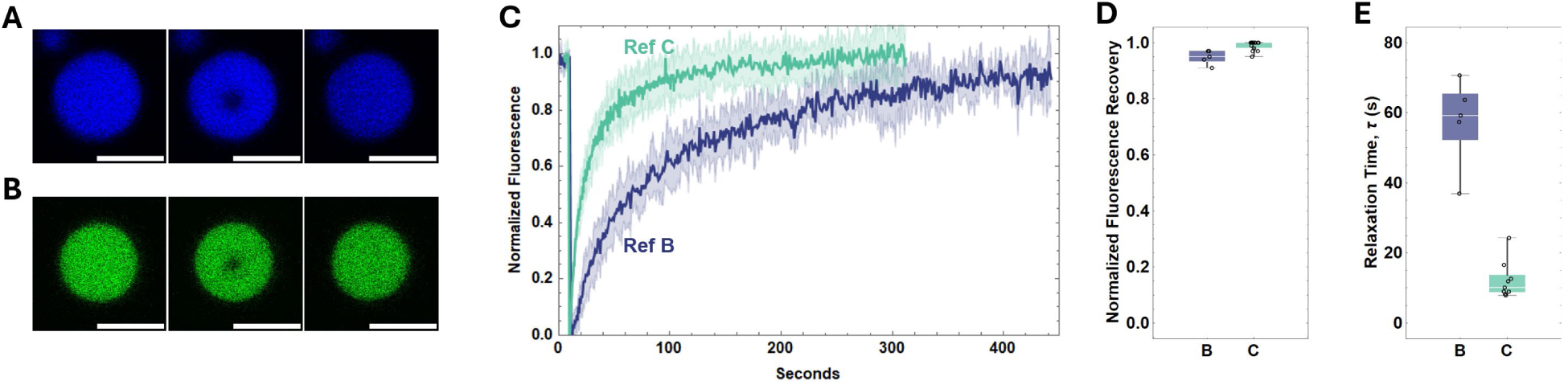
Diffusivity of reflectin condensates characterized by FRAP. FRAP of reflectin B and C condensates in 250 mM NaCl at pH 6 (MES, 25 mM). Fluorescence intensity of photobleached regions were corrected for incidental photobleaching, normalized, and fit to an equation of exponential decay to determine the fluorescence recovery and characteristic relaxation time, tau. Characteristic images of condensates of (A) reflectin B and (B) reflectin C pre-bleaching, post-bleaching and final. C) Normalized fluorescence intensity as a function of time for reflectin B (blue) and reflectin C (green). D) Normalized average fluorescence recovery and (E) average tau for reflectin B and reflectin C. Error bands in FRAP plots represent +/- 1 SD. Box and whisker plots display the median (center line), second and third quartile (solid box) and complete range (black fences) of the data. N=5 for reflectin B and N=15 for reflectin C. All scale bars are 5 μm.

**Figure 9.**
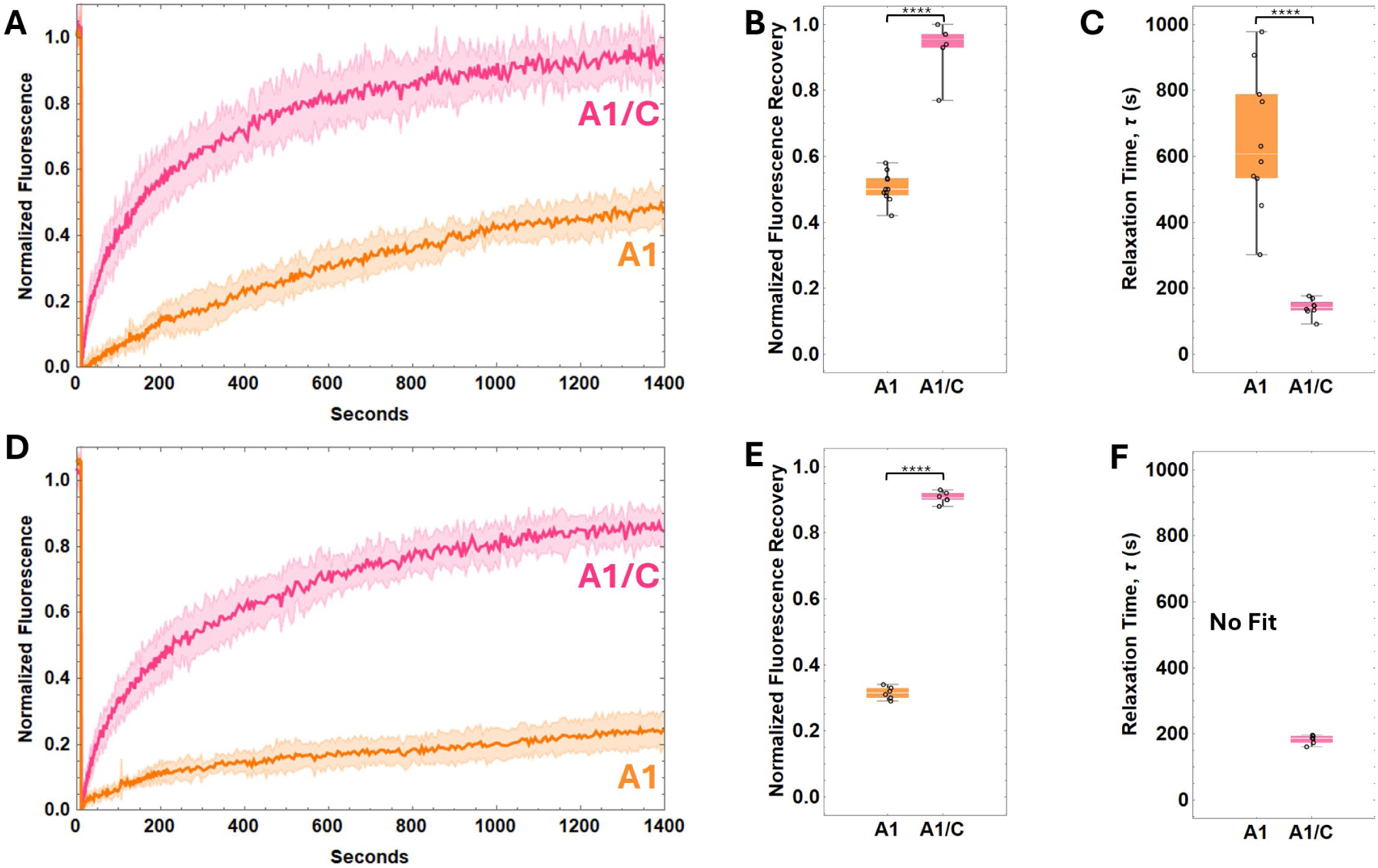
FRAP of reflectin A1 and A1/C condensates. As in Figure 8: (A) Normalized fluorescence recovery as a function of time, (B) total fluorescence recovery and (C) relaxation time tau for reflectin A1 and A1/C condensates in 250 mM NaCl at pH 6 (25 mM MES). (D) Normalized fluorescence recovery as a function of time, (E) total fluorescence recovery and (F) relaxation time tau for reflectin A1 and A1/C condensates in 250 mM NaCl at pH 7 (25 mM MOPS). Error bands in FRAP plots represent +/- 1 SD. Box and whisker plots display the median (center line), second and third quartile (solid box) and complete range (black fences) of the data. N=11 for reflectin A1 at pH 6, N=7 for reflectin A1 at pH 7, N= 8 for A1/C at pH 6 and N=6 for A1/C at pH 7. Data for A1 condensates previously published (23).

## Discussion

Cephalopod iridescence is mediated by periodically invaginated iridocyte cells, in which the alternating refractive indices of extracellular space and the reflectin-enriched, membrane-bounded lamellae form a distributed Bragg reflector that reflect light in an angle- and wavelength-dependent manner (6–8). Interestingly, dynamically tuned iridocytes regulated by neuronal control only have been found in the recently evolved Loliginid family of squids (9, 10). In these cells, acetylcholine reversibly activates reflectin phosphorylation which triggers reflectin condensation that in turn drives the proportional dehydration of the Bragg lamellae, shrinking their spacing and thickness and increasing their refractive index to tune the wavelength and intensity of reflectivity in a calibrated manner (5, 6, 11, 12). The protein NCD-responsive switch from A1 charge-stabilized assemblies to liquid condensates (23, 30) mirrors observed ultrastructural phase transitions from discrete 10 to 50 nm diameter particles to a uniform dense material upon phosphorylation of reflectin proteins observed *in vivo* (5, 9, 17). Further, isolated Bragg lamellae from acetylcholine-treated iridocytes from *Lolliguncula brevis* exhibited liquid characteristics such as surface wetting and fusion in marked contrast to the discrete puncta-filled lamellae isolated from untreated iridocytes (9).

The protein contents of neurotransmitter-tunable iridocytes from *Doryteuthis opalescens* are, by molarity, 11.6% A1, 10.5% A2, 36.1% B, and 41.9% C (6). Accordingly, we extended the *in vitro* model for phosphorylation-triggered LLPS of reflectin proteins to include A2, B and C in these proportions relative to A1. The enrichment of reflectins B and C in neurotransmitter-tunable iridocytes compared to non-tunable iridocytes (6), and their apparent uniqueness to species that possess neurotransmitter-tunable iridocytes (18) suggests they are critical for neuronally controlled LLPS within the Bragg lamellae. A previous DLS analysis of these ratios of reflectins A1, A2, B and C at a similar total protein concentration demonstrated coordinated assembly behavior (13). Individually, at pH 7 reflectin A1, A2, B and C formed soluble assemblies of approximately 12 nm r., 12 nm r., 70 nm r., and 100 nm r., respectively. Yet mixtures of these four proteins in the ratios found in either acetylcholine-responsive or non-responsive iridocytes formed soluble assemblies of approximately 15 nm r. with a similar size monodispersity to that of A1 alone, and narrower size monodispersity than that of A2, B, or C alone. This past observation of coordinated assembly in addition to the colocalization of reflectins A1, A2, B and C in assemblies shown in Figure 4C, F is evidence that hetero-oligomers are formed by these four reflectin proteins. Similarly, condensates formed from mixtures of A1/A2, A1/B and A1/C all displayed coordinated phase transitions for the range of protein NCDs and NaCl concentrations tested. In these two-protein condensates A2, B and C all inhibited the LLPS of A1 at high protein NCDs and either promoted (A2, C) or had little effect (B) on LLPS of A1 at low NCDs as a function of NaCl concentration. The liquid phase boundary of both neurotransmitter-tunable and non-tunable mixtures of A1/A2/B/C is dominated by reflectin B, as seen in the similarity between all three phase diagrams (**Figure 3G, 4E)**.

Within the Bragg lamellae of neurotransmitter-tunable iridocytes reflectins, A2, B and C could sharpen the phosphorylation-driven liquid phase transition by stabilizing the ‘off’ state for unphosphorylated (high protein-NCDs) reflectins A1 and A2 and promoting LLPS upon phosphorylation (low protein-NCDs). At high protein NCDs, A2, B and C could inhibit LLPS of A1 by a capping mechanism that arrests condensate growth (31–33). The increased numbers of amino acid residues capable of forming weak non-covalent interprotein interactions such as salt bridges or arginine-tyrosine cation-ϖ bond, has been observed to increase the thermodynamic drive for a protein to undergo LLPS (24, 34–39). Of the reflectins, A1 and A2 have the greatest number of arginine and tyrosine residues (38R: 44Y and 27R:29Y respectively), whereas B and C have the least (27R:15Y and 18R:11Y respectively). At protein NCDs and NaCl concentrations where reflectin A1 normally exists as two phases, heterotypic A1/A2, A1/B, or A1/C interactions may out-compete homotypic A1/A1 interactions and inhibit LLPS of A1 by exhausting A1’s valencies (32, 36). Once the protein-NCD is lowered or the NaCl concentration to conditions in which A2, B and C each exists as two phases, homotypic A2/A2, B/B and C/C interactions lessen the competition between heterotypic A1/A2, A1/B and A1/C interactions and homotypic A1/A1 interactions, increasing the effective valency of A1 and allowing LLPS of A1. Mutational analyses that skew the Arg:Tyr ratio of A1 could test the hypothesis that the uneven Arg:Tyr ratios in B and C could further act to restrict condensate growth and LLPS (31, 32).

We previously hypothesized that LLPS of the Bragg lamellar contents would provide an environment conducive to the dynamic enzymatic phosphorylation and dephosphorylation of reflectins (23) while maintaining the high protein density necessary to maximize both the reflectivity and the osmotic tunability of the Bragg lamellae. The protein concentration within the Bragg lamellae was previously measured to be 381 mg/ml (15), and determined to be 416 +/- 42 mg/ml for reflectin A1 condensates (23). Although the FRAP recovery of reflectin A1 condensates at physiologically relevant conditions was slow and incomplete (23), our present findings show that diffusivity is increased in reflectin B and reflectin C condensates, and that reflectin C greatly increases the diffusivity of A1 in A1/C condensates. This suggests that reflectins B and C could facilitate both the signal cascade that culminates in reflectin phosphorylation (15) and constitutively expressed phosphatases by increasing the rates of diffusion of the reflectins and their respective kinases and phosphatases and thus the rates of these reactions (40, 41).

Reflectin protein distribution within the Bragg lamellae was previously observed to be spatially organized, with reflectin C localized at the membrane edges, and reflectins A1, A2 and B colocalized in the lumens of the lamellae (6). Our observations of the multiphasic spatial organization of reflectin condensates can be explained by partial miscibility of the different, individual reflectin condensates, and this in turn can explain the inhomogeneous spatial organization of the reflectins previously observed within the Bragg lamellae *in vivo* (6). Further, a recently reported analysis demonstrates that this inhomogeneous spatial segregation can create the gradient of refractive index across the Bragg lamella that is essential for maximal tunability of the Bragg reflector in *Doryteuthis* squids (42).

Condensates formed from the mixture of reflectins A1, A2, B and C in the ratio found in Bragg lamellae of the neurotransmitter-tunable iridocytes in *Doryteuthis opalascens* drastically changed their multi-component organization in response to charge-neutralization (decreased NCD) of the reflectins. At high protein NCDs, A1 and A2 were largely compartmentalized within the surrounding B or C of the condensates, and this organization reversed at low protein NCDs (**Figures 6D,E,F and 7A, E, I)**. These results suggest that the surface tension of reflectins A1 and A2 is higher than B and C at high protein NCDs and lower at low protein NCDs. Significantly, the organization of reflectins in condensates formed from a mixture of A1, A2, B and C in the ratio found in non-tunable iridocytes showed no such organization (**Figures 6A, B, C and 7B)**. Presumably, this organization is controlled both by the relative surface tensions of each compartment (28, 43) and the relative hydrophobicities and resulting co-solubilities of A1, A2, B and C. Segregation within the multicompartment condensates was never complete, and the condensate architecture did not minimize the interfacial area between compartments (**Figures 6D, E, F, and 7)**. At low protein NCDs (corresponding to the state of phosphorylation-driven charge neutralization) reflectins B and C were consistently partitioned to the droplet interior (**Figures 6D and 7A, E, I)** and the reverse was true for high protein NCDs (corresponding to the unphosphorylated state) (**Figure 6E, F)**, suggesting that the relative interfacial energies between reflectins A1, A2, B and C and solvent dictated the multicompartment organization and these interfacial energies changed with protein NCD, as they would with neurotransmitter-activated phosphorylation. These results are consistent with the proteins adopting different conformations and thus presenting different surfaces with different hydrophobicities to solvent as a function of protein NCD, as has previously observed for reflectin A1 assemblies (13, 20). However, no such analyses have been performed for reflectin liquid condensates.

While we investigated LLPS of reflectins A1, A2, B and C in the absence of a membrane, the interfacial preferences revealed by the dynamic multiphase condensates can be extended to interactions with the Bragg lamellar membrane. For neurotransmitter-responsive mixtures of reflectins A1, A2, B and C, we observed condensates with partial immiscibility of reflectins A1 and A2 with reflectins C and B (**Figures 6D,E,F, 7A)** with A1 and A2 enveloping B and C. *In vivo,* the observed spatial segregation described above can be understood as the partial immiscibility of reflectins A1, A2 and B with C, with C enveloping A1, A2, and B. The presence of the lamellar membrane would present an interface much more hydrophobic then the aqueous solvent used in our in vitro analyses, and could thus invert the spatial organization of the multicompartment condensates observed *in vitro*. Furthermore, reflectin C attenuated the rearrangement of giant unilamellar vesicles by reflectin A1 (44), further demonstrating the lipid membrane affinity of reflectin C. The NCD dependence of the condensate’s external interface would enable changing the energetics of protein-membrane interactions depending on the phosphorylation state of the reflectins, possibly contributing to maintenance of the regularity of Bragg lamellar morphology through its cyclical dehydration and rehydration.

The dynamic multicompartment condensates of reflectins A1, A2, B and C provide the potential to tunably provide reflectins A1 and A2 with a separate chemical environment from B and C. This could facilitate differential phosphorylation of the spatially segregated reflectins by kinases and phosphatases that we assume are colocalized with reflectin proteins in the lamella due to the rapid reversible dehydration of the lamella One difficulty posed to *in vitro* models is that reflectins A1 and A2 are phosphorylated at Y14, Y127, H118 and Y12, Y214, S218, Y223, respectively, upon iridocyte activation while reflectin B is partially dephosphorylated at S228 and S259, and is constitutively phosphorylated at T63 and S228. (6, 12). A1 and A2 have a similar phosphorylation motif that differs from that of B (6), and the segregation of A1 and A2 from B could differentially regulate kinase and phosphatase activities in the Bragg lamellae as has been shown for other protein condensates (45–47). Neurotransmitter-tunable iridocytes are constitutively ‘off’ until activated by ACh; removal of ACh causes the Bragg lamellae to rehydrate and the wavelength of reflected light to increase (15). The spatial segregation of A1 and A2 from B upon decreasing protein NCD (as occurs upon phosphorylation) may thus be an integral response to the ACh-triggered signal cascade that drives Bragg lamellar dehydration and the resulting change in color.

The reliance of the squid’s remarkable reflectin-controlled biophotonic system on phase changes in soft living matter has inspired numerous biomimetic applications for reflectin proteins (48–53) as well as methods for controlling reflectin assembly (20–22). We present a significant extension of *in vitro* reflectin material tunability by introducing protein-specific effects on phase transitions and liquid condensate properties as well as protein-NCD dependent spatial organization of different reflectin protein species in dynamically tunable liquid condensates. We relate these findings to their potential for enhancing the *in vivo* control of cyclable dehydration/rehydration and reflectivity of the Bragg lamellae in which they are contained. Several mechanisms of this control are identified, including: (1) increased sensitization to phosphorylation-triggered LLPS by reflectins A2, B and C; (2) increased diffusivity within the dense phase by reflectins B and C, consequently increasing the rates of reaction to more rapidly tuning the instantaneous balance between phosphorylation and dephosphorylation; and (3) dynamic spatial regulation of signal cascade components; and (4) dynamically switching reflectin-membrane interactions. Further investigations of the phosphorylation- and NCD-controlled changes in properties (including surface tension, hydrophobicity, and interactions with membranes) of these reflectin liquid condensates will help elucidate the thermodynamic drivers of their dynamic multiphase organization and advance the engineering of reflectin-based biomaterials.

## Experimental procedures

### Protein expression and purification

Reflectin A1, A2, B and C were purified as previously described (13, 20, 23). Codon-optimized sequences of reflectins A1, A2, B and C were cloned into pj411 plasmids. Proteins were expressed in Rosetta 2 (DE3) cells grown in in 1 L (A2, B,C) or 2 L (A1) cultures at 37 ° C from plated and sequenced transformants in the presence of 50 mg/mL kanamycin. Expression was induced at an absorbance of 0.6 to 0.7 with 5 mM IPTG. After 16 hours, cell cultures were pelleted by centrifugation and frozen at −80 ° C. Reflectins were expressed in inclusion bodies which were purified using BugBuster medium (Novagen, Inc) per manufacturer protocol, then resolubilized in 8 M urea 5% acetic acid. Reflectins were purified using a 10 ml Hitrap cation-exchange column (Cytiva) and eluted using a step gradient (from 5% acetic acid, 8 M urea to 7 then 10% 5% acetic acid, 6 M guanidinium chloride for reflectins A1 and A2, and 0-5-7% 5% acetic acid, 6 M guanidinium chloride for B and C). Purity of collected fractions was determined by SDS-PAGE and A260/A280 (0.55-0.57), which were pooled and loaded onto a reverse-phase HPLC XBridge 4.6 ml C4 column (Waters) equilibrated with 10% acetonitrile with 0.1% TFA and 90% water with 0.1% TFA (A1 and A2) or 5% acetonitrile with 0.1% TFA and 95% water with 0.1% TFA (B and C) and eluted over a gradient to 100% acetonitrile with 0.1% TFA. Fractions were frozen at −80 ° C or shell frozen using an ethanol and dry ice bath, lyophilized, and stored at −80 ° C. Lyophilized reflectin was solubilized using 0.22 μm-filtered acetic acid buffer (pH 4, 25 mM) and dialyzed using two 12 hour changes of 1000X sample volume of the same buffer at 4° C. Protein concentration was calculated using absorbance at 280 nm and molar absorption coefficients for each protein (reflectin A1: 120 685 M^-1^ cm^-1^, reflectin A2: 76335 M^-1^ cm^-1^, reflectin B: 22350 M^-1^ cm^-1^, reflectin C: 19458 M^-1^ cm^-1^). Protein stocks were centrifuged for clarification of monomers at 18,000g for 15 min at 4° C immediately prior to use in all assays. Protein stocks were stored at 4° C between uses.

### Fluorescent labeling

Reflectins A1-C232S (23, 54) and C were covalently labeled by conjugation with cysteine-specific fluorescein-methanethiosulfonate (FMTS) and reflectin A2 with sulforhodamine-methanethiosulfonate (RMTS). 50 μM protein stocks were incubated with 500 μM FMTS or RMTS in acetic acid buffer (pH 4, 25 mM) with gentle orbital stirring for 4 h at room temperature then overnight at 4° C. Labeled reflectins were concentrated using 10 KDa molecular weight cut off spin filters (Amicon) and excess label removed using the same HPLC and storage method as for initial protein purification. Reflectins B and C were covalently labeled using maleimide-based Alexafluor-405 (reflectin B) and Alexafluor-660 (reflectin C). To ensure reflectins B and C remained monomeric in pH 7 labeling conditions, protein was dialyzed into 8 M urea solution containing acetic acid buffer (pH 4, 25 mM) using one 24 h exchange of 1000X sample volume. Protein was then dialyzed into 8 M urea solution containing MOPS buffer (pH 7, 25 mM). 50 μM protein stocks were incubated with 500 μM Alexafluor-405 or −660 with gentle orbital stirring for 4 h at room temperature then overnight at 4° C. Samples then were dialyzed into 8 M urea solution containing acetic acid buffer (pH 4, 25 mM) and then into acetic acid buffer (pH 4, 25 mM), concentrated using10 KDa molecular weight cut off spin filters (Amicon) and any remaining excess label removed using the same HPLC and storage method as described for the initial protein purification. All labeling efficiencies were measured and confirmed to be approximately 100%.

### Phase diagrams and droplet imaging

For determination of phase diagrams, unlabeled reflectins respectively contained 5% fluorescein-labeled reflectin A1 C232S, 1% sulforhodamine-labeled A2, 0.5% Alexafluor-405-labeled reflectin B, and 0.5% Alexafluor-660-labeled reflectin C. To determine the liquid phase boundaries, 0.6 uL of 100 µM protein in acetic acid buffer (pH 4, 25 mM) were diluted into 9.4 µl of experimental buffer to a final protein concentration of 4 µM. For imaging of multiphase condensates, 0.6 µL of 200 µM protein in acetic acid buffer (pH 4, 25 mM) were diluted into 9.4 µl of experimental buffer to a final protein concentration of 4 µM. Solutions were mixed gently, and incubated for 10 min at 25 ° C for 10 min using 0.5 ml protein LoBind tubes (Eppendorf). The solutions were pipetted onto cleaned glass coverslips and imaged immediately after deposition using a Leica SP8 resonant confocal microscope with 63X objective (NRI-MCDB Microscopy Facility). Multiphase droplets were imaged similarly but using PEG-passivated chamber slides which were sealed with fast-drying clear nail polish. All buffers were freshly 0.22 μm-filtered before use. NaCl solutions were buffered with 25 mM acetic acid buffer for pH values of 4, 4.5, 5 and 5.5; 25 mM MES buffer for pH values of 6 and 6.5; and 25 mM MOPS buffer for pH values of 7, 7.5 and 8. Ex_max_ and Em_max_ for the fluorophores used are: Alexafluor 405 405 nm and 425 nm; fluorescein 488 nm and 518 nm; sulforhodamine 578 nm and 590 nm; Alexafluor 660 600nm and 690 nm.

### FRAP experiments and analysis

Protein stocks for single-protein condensate experiments were 100 μM of reflectin A1 containing 5% fluorescein-labeled A1 C232S or reflectin C containing 5% fluorescein-labeled reflectin C; protein stocks for A1/C condensates were 100 μM of the indicated mole percent A1 containing 5% fluorescein-labeled A1 C232S and the indicated mole percent reflectin C. Protein stocks were diluted into freshly 0.22 μm-filtered buffers and then immediately pipetted into PEG-passivated chamber slides which were sealed with fast-drying clear nail polish. Using the FRAP module of the Leica SP8 scanning confocal microscope, a 1 μm diameter circular ROI [region of interest; Point ROI” in LASX (https://www.leica-microsystems.com/products/microscope-software/p/leica-las-x-ls/) software] was selectively photobleached at 45 to 50% laser power for 150 to 200 ms. The bleached ROI was monitored for 10 s before bleaching and for 300 s to 25 m post-bleaching. For data analysis, videos were stabilized in ImageJ (https://imagej.net/ij/) using the StackReg plugin (https://bigwww.epfl.ch/thevenaz/stackreg/) if necessary, and the fluorescence intensity of the bleached region, an unbleached region in the droplet, and background were used to correct for photobleaching using the equation:

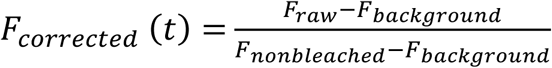

where F_raw_ is the fluorescence intensity of the phobleached region, F_background_ the fluorescence intensity of the background and F_nonbleached_ the fluorescence intensity of the ROI pre-bleaching. Data were fully normalized using the equation:

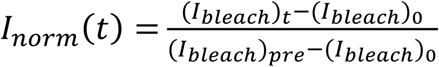

where (I*_bleach_*)*_t_* is the intensity of the photobleached region at time t, (I*_bleach_*)_0_ is the intensity of the photobleached region at the time of photobleaching, and (I*_bleach_*)*_pre_* is the intensity of the photobleached region prior to photobleaching. To determine the characteristic relaxation time *τ* data were fit to the equation:

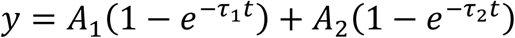

using the NonLinearRegression function in Mathematica.

## Acknowledgements

Research was sponsored by the U.S. Army Research Office and accomplished under cooperative agreement W911NF-19-2-0026 for the Institute for Collaborative Biotechnologies. The content of the information does not necessarily reflect the position or the policy of the Government, and now official endorsement should be inferred. We thank Ben Lopez of UCSB’s NRI-MCDB Microscopy Facility for his expert assistance with laser confocal microscopy

