## Supplemental Information for "Reflectins form multicompartment liquid-liquid phase separated condensates that mirror and may facilitate spatial organization in squid skin Bragg lamellae"

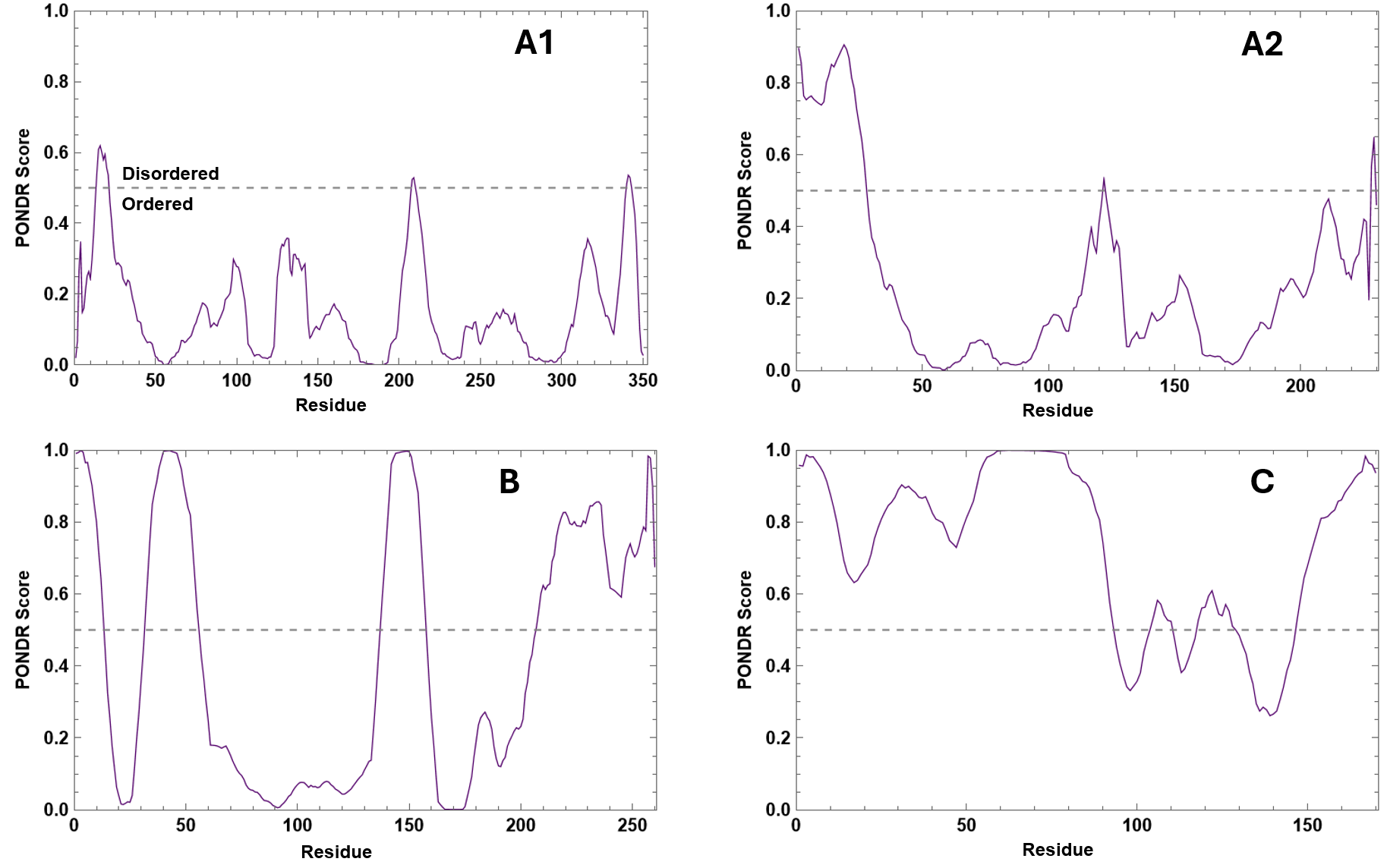

Figure S1. Predictor of Natural Disordered Regions (PONDR) for reflectins A1, A2, B and C. PONDR scores of individual amino acids for each of the four reflectin proteins in this study determined using a sliding window of 9 amino acids. A residue is considered disordered if its value matches or exceeds 0.5. Average prediction scores are 0.1552 for A1; 0.2432 for A2; 0.4223 for B; and 0.7295 for C.

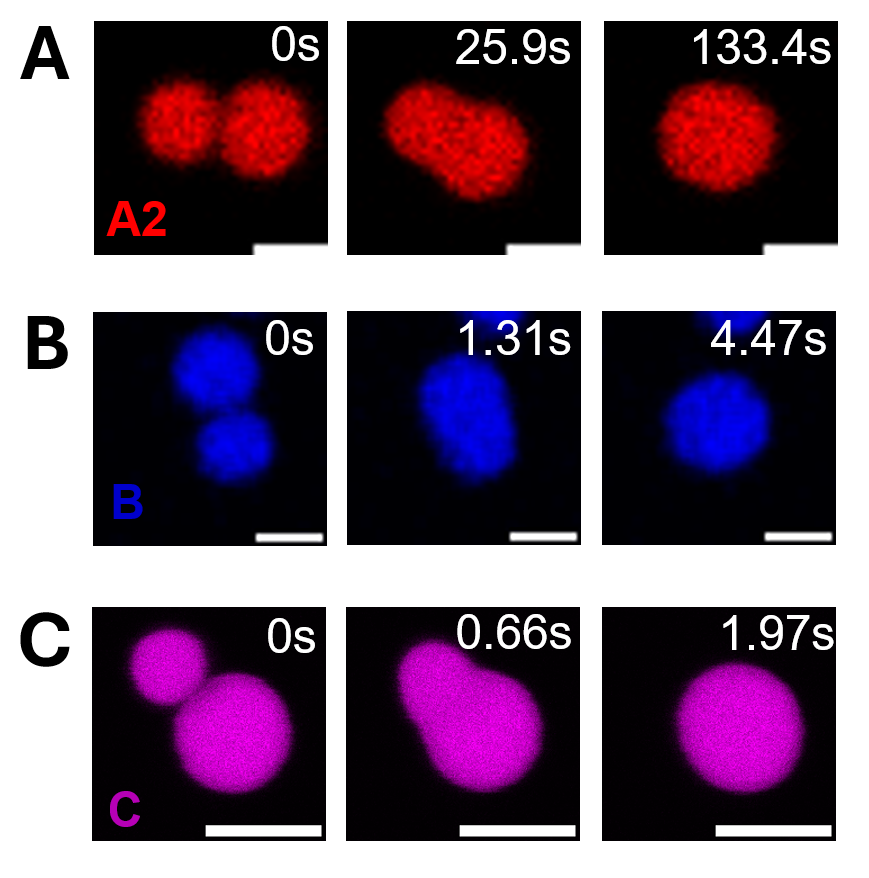

Figure S1. Reflectin liquid-like droplets fuse and relax to sphericity. Time lapses of liquid-like droplets of reflectins A2 (A), B (B) and (C) fusing and relaxing to sphericity. Scale bars are 5 μm.

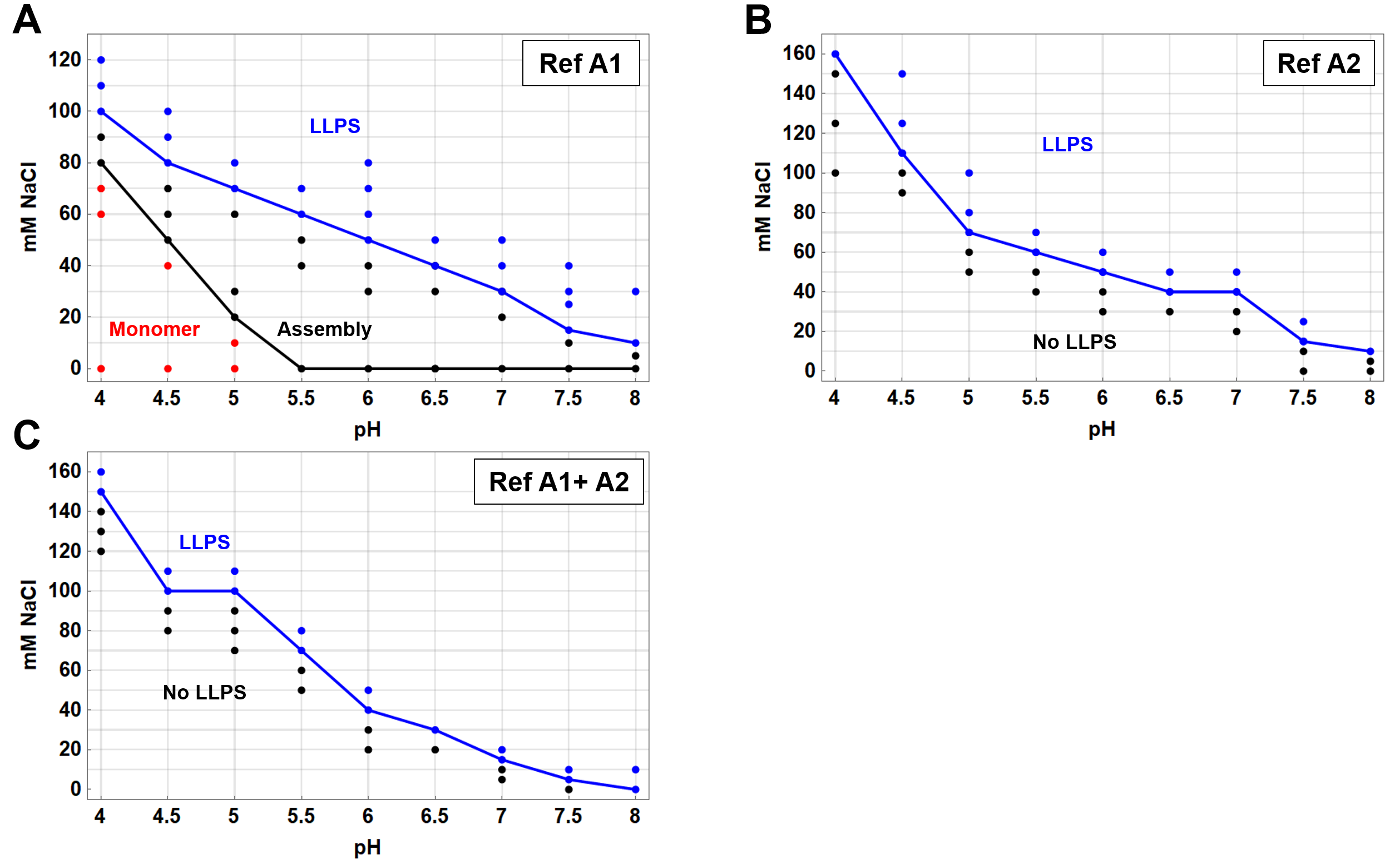

Figure S2. Phase diagrams of reflectin A, A2, and A1/A2 as a function of NaCl concentration and pH. A) One hundred μM partially fluorescently labeled reflectin A1 (see methods: protein labeling), (B) reflectin A2 and (C) a solution of 52.6 and 47.37 μM of partially fluorescently labeled A1 and A2 respectively was diluted to a final protein concentration of 4 μM. In (A), red dots represent monomer as detected by DLS, black dots represent assemblies detected either by DLS or confocal microscopy, and blue dots represent detection of liquid droplets by confocal microscopy. For B and C, black dots represent lack of detection of liquid droplets and blue dots represent detection of liquid droplets by confocal microscopy.

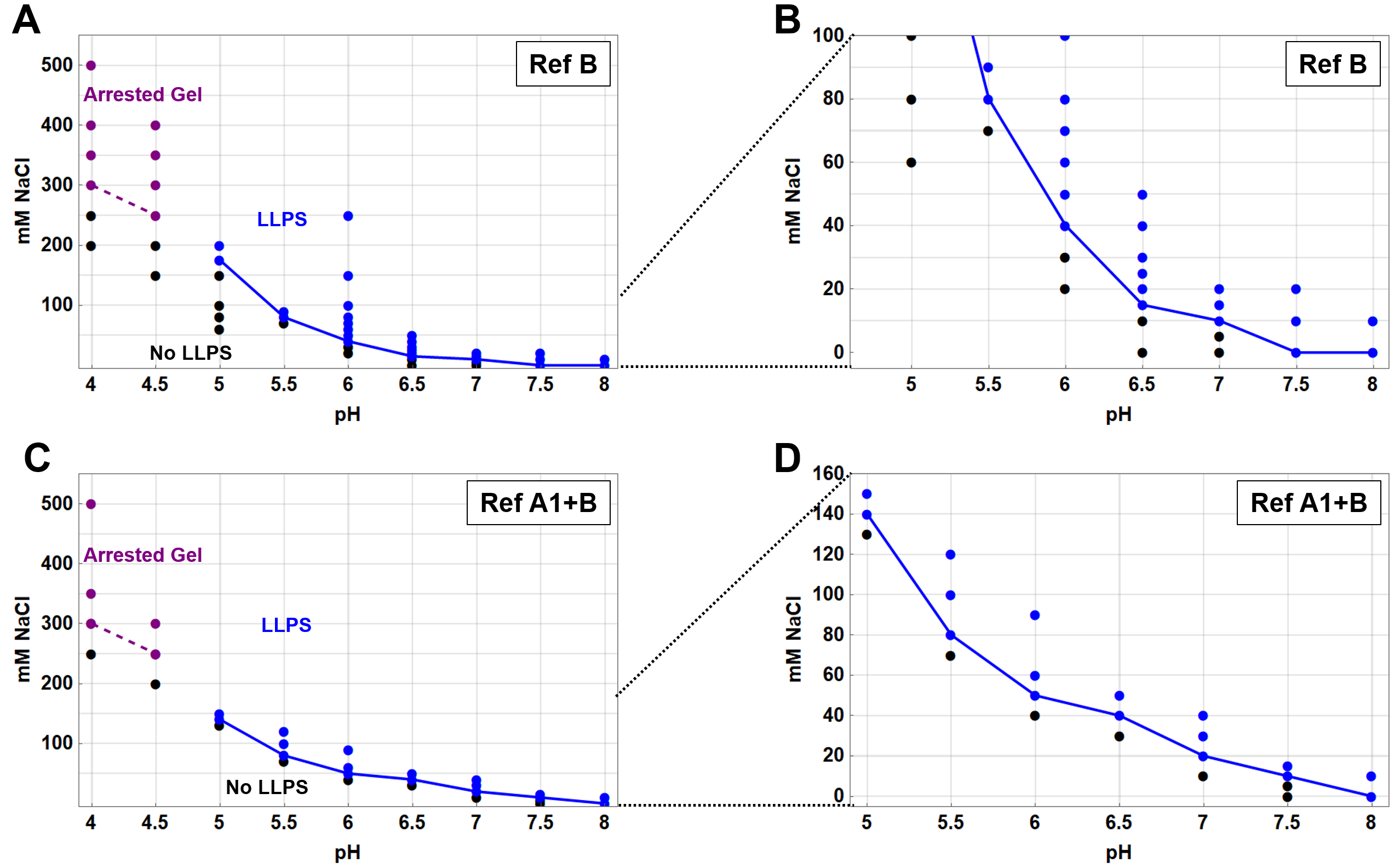

Figure S3. Phase diagrams of reflectin B and A1/B as a function of NaCl concentration and pH. A,B) One hundred μM partially fluorescently labeled reflectin B (see methods: protein labeling) or (B,D) a solution of 24.4 and 75.6 μM of partially fluorescently labeled A1 and B respectively was diluted to a final protein concentration of 4 μM. Purple dots represent arrested gel-like condensates, black dots represent lack of detection of liquid droplets or arrested gel-like condensates and blue dots represent detection of liquid droplets by confocal microscopy.

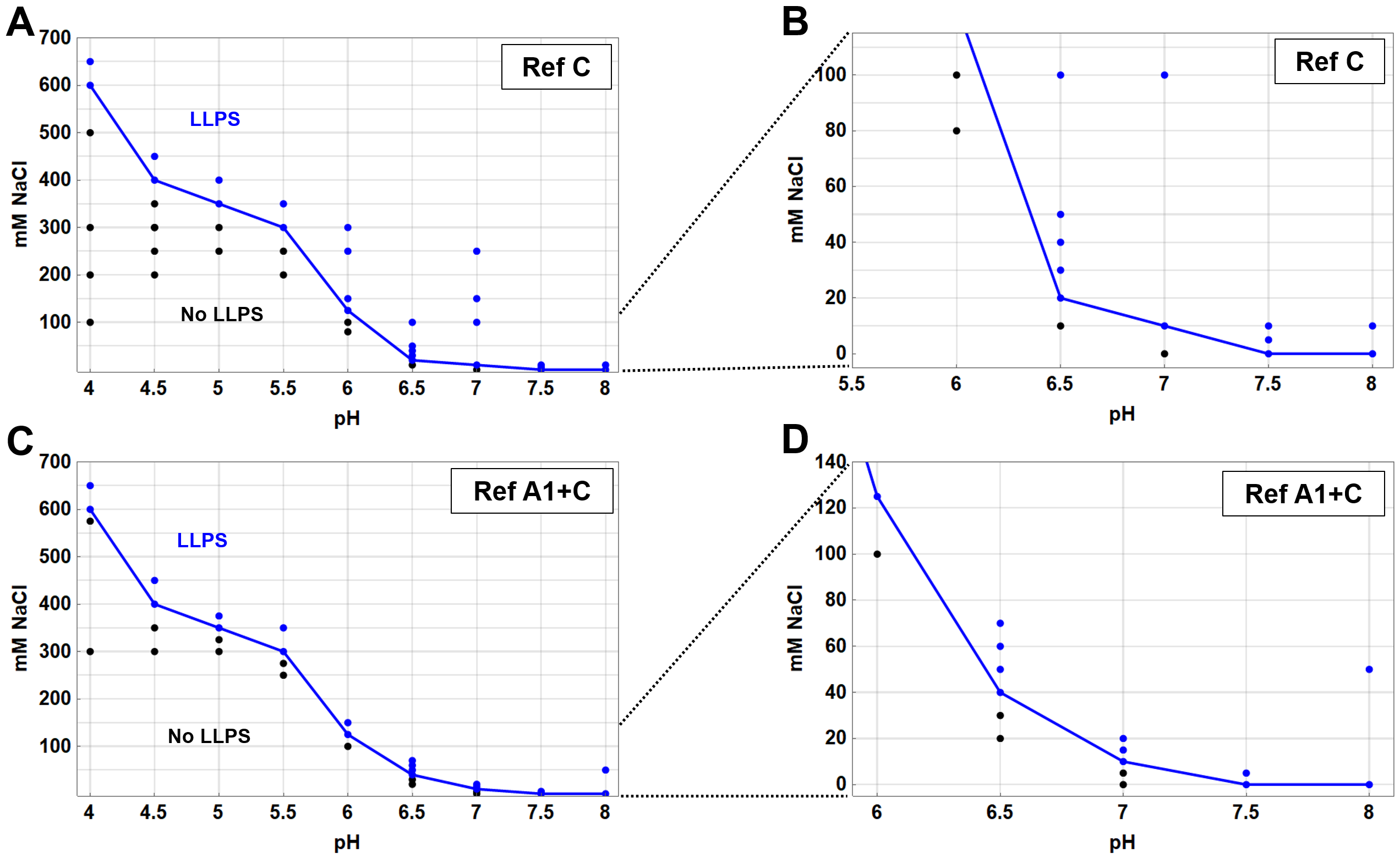

Figure S4. Phase diagrams of reflectin C and A1/B as a function of NaCl concentration and pH. A,B) One hundred μM partially fluorescently labeled reflectin C (see methods: protein labeling) or (B,D) a solution of 21.7 and 78.6 μM of partially fluorescently labeled A1 and C respectively was diluted to a final protein concentration of 4 μM. Black dots represent lack of detection of liquid droplets and blue dots represent detection of liquid droplets by confocal microscopy.

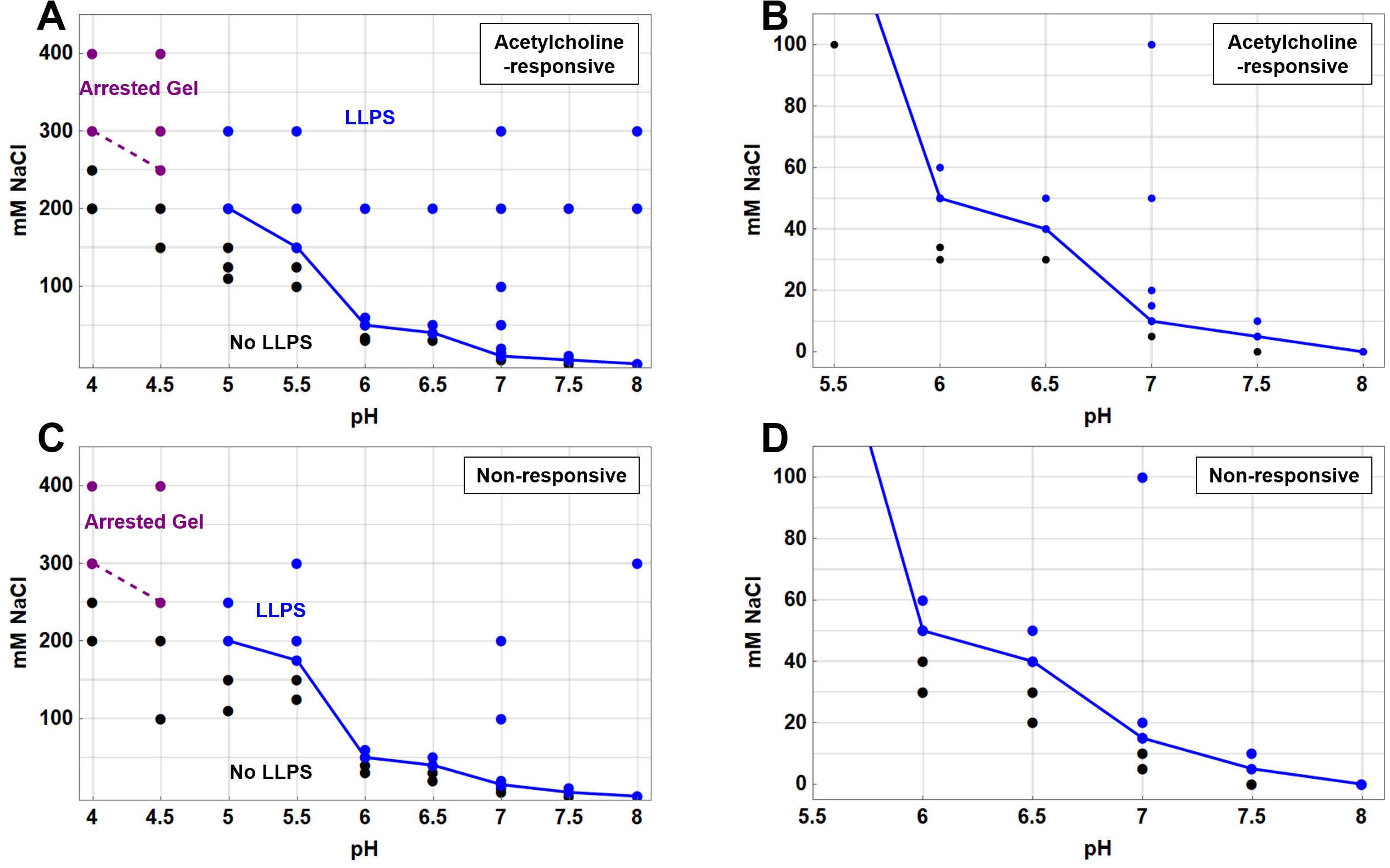

Figure S5. Phase diagrams of molar ratios of reflectins found in acetylcholine- and non-responsive iridocytes as a function of NaCl concentration and pH. A) Liquid phase boundary (blue) of acetylcholine-responsive mixture and (B) inset of same plot. C) Liquid phase boundary (blue) of non-responsive mixture and (D) inset of same plot. Purple dots represent arrested gel-like condensates, black dots represent lack of detection of liquid droplets or arrested gel-like condensates and blue dots represent detection of liquid droplets by confocal microscopy.

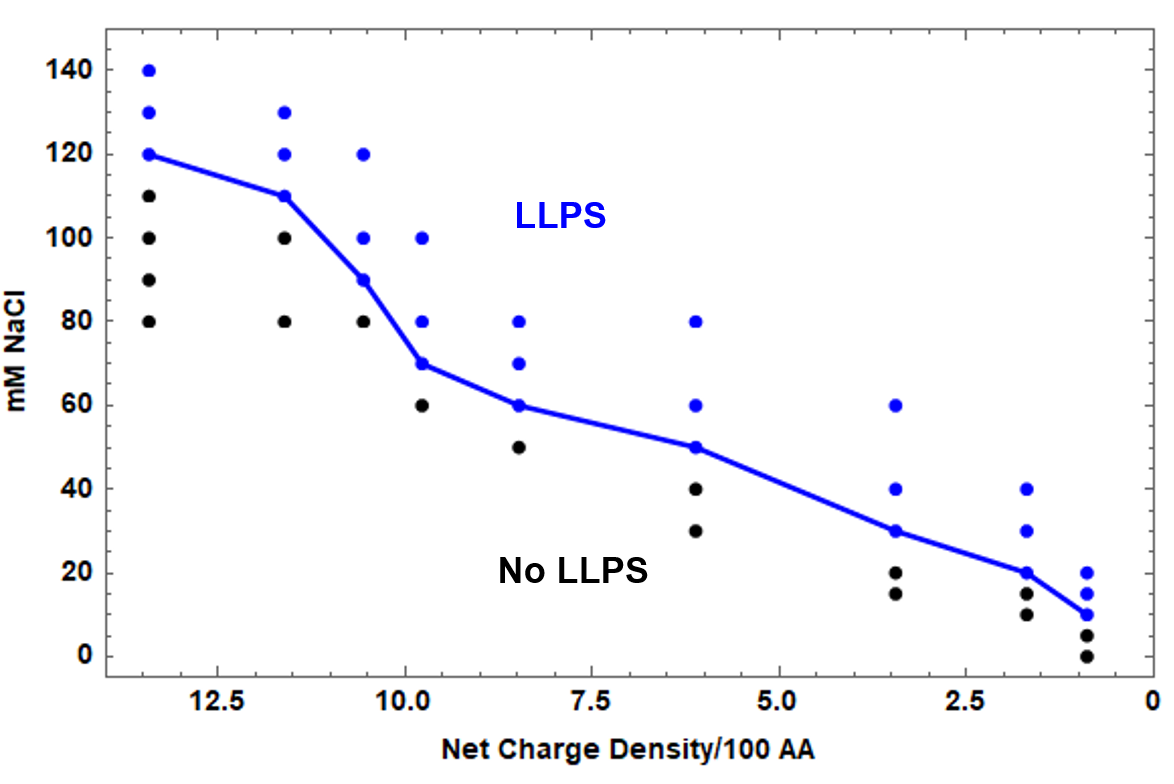

Figure S6. Liquid phase boundary of reflectin A1 with total protein concentration of 0.5 μM as a function of calculated protein net charge density and NaCl concentration. Liquid condensates (blue) were distinguished from the absence of liquid condensates (black) by confocal microscopy. Black dots represent lack of detection of liquid droplets and blue dots represent detection of liquid droplets by confocal microscopy.
